# Tumor-expressed SPPL3 supports innate anti-tumor immune responses

**DOI:** 10.1101/2024.04.25.591102

**Authors:** Tamara Verkerk, Antonius A. de Waard, Sofie J.I. Koomen, Jasper Sanders, Tineke Jorritsma, Anouk T. Pappot, Nordin D. Zandhuis, Tao Zhang, Manfred Wuhrer, Hannes S.J. Stockinger, Klaas P.J.M. van Gisbergen, Robbert M. Spaapen, S. Marieke van Ham

## Abstract

The development of an effective anti-tumor response relies on the synergistic actions of various immune cells that recognize tumor cells via distinct receptors. Tumors, however, often manipulate receptor-ligand interactions to evade recognition by the immune system. Recently, we highlighted the role of neolacto-series glycosphingolipids (nsGSLs), produced by the enzyme β1,3-*N*-acetylglucosaminyltransferase 5 (B3GNT5), in tumor immune escape. We previously demonstrated that loss of signal peptidase like 3 (SPPL3), an inhibitor of B3GNT5, results in elevated levels of nsGSLs and impairs CD8 T cell activation. The impact of loss of SPPL3 and an elevated nsGSL profile in tumor cells on innate immune recognition remains to be elucidated. This study investigates the anti-tumor efficacy of neutrophils, NK cells, and γδ T cells on tumor cells lacking SPPL3. Our findings demonstrate that SPPL3-deficient target cells are less susceptible to trogocytosis by neutrophils and killing by NK cells and γδ T cells. Mechanistically, SPPL3 influences trogocytosis and γδ T cell instigated killing through modulation of nsGSL expression while SPPL3-mediated reduced killing by NK cells is nsGSL-independent. The nsGSL-dependent SPPL3 sensitivity depends on the proximity of surface receptor domains to the cell membrane and the affinity of receptor-ligand interactions as shown with various sets of defined antibodies.

Thus, SPPL3 expression by tumor cells alters crosstalk with immune cells through the receptor-ligand interactome thereby driving escape not only from adaptive but also from innate immunity. These data underline the importance of investigating a potential synergism of GSL synthesis inhibitors with current immune cell activating immunotherapies.

## Introduction

To establish an effective anti-tumor response, immune cell receptors work synergistically to mediate and fine-tune cellular responses. Functional interaction between receptors and their ligands is necessary to activate the right pro-inflammatory immune cells or inhibit suppressive cells for a specific, yet balanced immune response. Well-known is the human leukocyte antigen class I (HLA-I) which presents (neo)antigens from an infected or mutated cell, which are recognized by cognate T cell receptors (TCRs) on CD8 T cells, instigating T cell activation and anti-tumor effector function upon interaction^1^. Unfortunately, pro-inflammatory receptor-ligand interactions can be impeded or modulated by tumor cells to avoid proper recognition by the innate and adaptive immune system. Tumor evasion strategies have extensively been studied and new tumor evasion strategies focusing on tumor-specific traits are still being discovered.

Recently, we demonstrated an important role for signal peptidase like 3 (SPPL3) in supporting adaptive immune responses through inhibition of the neolacto-series glycosphingolipid (nsGSL) synthesis pathway. We identified that SPPL3, in addition to the previously described destruction of enzymes involved in *N*-glycosylation of proteins, also destroys the enzyme β1,3-*N*-acetylglucosaminyltransferase 5 (B3GNT5), which is a key catalyst in the production of nsGSLs^8–10^. In general, GSLs are fundamental components of the cell membrane which regulate cell proliferation, differentiation, protein function and intercellular receptor-ligand interactions in health, but may also contribute to pathology^2–7^. Our data showed that elevated B3GNT5-produced nsGSL levels impair accessibility of HLA-I leading to a reduced CD8 T cell activation *in vitro*^11^. In addition, we showed that antibody binding to HLA-I domains proximal to the plasma membrane is relatively more affected by nsGSLs than to HLA-I regions that were more distal to the plasma membrane. Moreover, the interactions of LIR-1 and killer cell immunoglobulin-like-receptor-2DL (KIR2DL2) fusion proteins with HLA-I were impaired by nsGSLs, implying that nsGSLs can interfere important immune receptor-ligand interactions.

Another study demonstrated that CD19 on SPPL3-negative target cells was badly bound by antibodies leading to increased resistance to CD19 CAR T cell activity. In this case, SPPL3 directed this effect through control of *N*-glycosylation of CD19^12^.

There are also studies which implicate that loss of SPPL3 may impair innate immune responses. One study showed that suppression of nsGSL synthesis through SPPL3 activity allows for better accessibility of the complement regulator CD59^13^. Another study illustrated that genetic loss of SPPL3 led to a decreased susceptibility of B cell lymphoma cell line NALM6 to NK cell mediated cytotoxicity^14^. These findings were also observed by Zhang and colleagues who, additionally, showed impaired binding of NK cell receptors NKG2D and CD2 to their partners MICA/B and CD58 on cells lacking SPPL3^15^. Yet, whether the underlying mechanism of these observations is through modulation of GSL or protein glycosylation is not clear.

Together with our findings, this implies that the regulatory function of SPPL3 may control both adaptive and innate immune responses, possibly through nsGSLs and *N*-glycosylation. On a molecular level, this may be regulated through determining accessibility of a potentially large variety of membrane proteins.

In this study, the nsGSL-dependent and -independent role of regulation by SPPL3 in target cells on neutrophil, NK cell and γδ T cell function was first explored. We show that target cells lacking SPPL3 are protected against trogocytosis by neutrophils and killing by NK cells and γδ T cells. Mechanistically, modulation of nsGSL expression by SPPL3 formed the basis for the SPPL3-sensitivy of trogocytosis by neutrophils or killing by γδ T cells, but not of killing by NK cells. We then determined SPPL3- and nsGSL-sensitivity for many individual cell surface receptors, and established that membrane receptor size and affinity between interacting proteins are crucial for nsGSL driven shielding. Interestingly, where most receptor-ligand/antibody interactions were SPPL3-sensitive through nsGSL modulation, some interactions were affected by SPPL3 in an nsGSL-independent fashion. All together, these results show a new mode of regulation of innate immune responses. Upon decreased expression or loss of SPPL3 by tumor cells, the cumulative immune escape effect may therefore involve both adaptive and innate anti-tumor immunity.

## Materials and Methods

All antibodies and membrane dyes used are listed in *table 1 and 2*.

**Table 1:** Overview of antibodies and membrane dyes used for flow cytometry.

| Antibody | Clone | Conjugate | Dilution or concentration | Supplier | Reference |
| --- | --- | --- | --- | --- | --- |
| Anti-human TCR Vδ2 | B6 | PE | 1:200 | BioLegend | 331408 |
| Anti-human CD16 | 3G8 | BV605 | 1:500 | BioLegend | 302040 |
| Anti-human CD56 | NCAM16.2 | BV510 | 1:100 | BD Horizon | 563041 |
| Anti-human CD3 | UCHT1 | BV510 | 1:200 | BD Horizon | 563109 |
| Anti-human CD25 | 2A3 | BV605 | 1:100 | BD Horizon | 562660 |
| Anti-human CD69 | FN50 | BUV395 | 1:100 | BD Biosciences | 564364 |
| Anti-human/mouse Granzyme B | GB11 | FITC | 1:100 | BD Biosciences | 560211 |
| Anti-HLA-I/MHC-I | W6/32 | PerCP-eFluor 710 | 1:250 | Invitrogen | 46-9983-42 |
| Anti-human CD11b | Bear1 | PE | 1:100 | Beckman Coulter | IM2581U |
| Anti-human ICAM1 | 15.2 | AF546 | 1:100 | Santa Cruz | Sc-107<br>AF546 |
| Goat anti mouse IgG | polyclonal | AF647 | 1:200 | Invitrogen | A21236 |
| Goat anti mouse IgG | M1310G05 | APC | 1:200 | Biologend | 410712 |
| LIVE/DEAD<sup>TM</sup> Fixable Near-IR |  |  | 1:800 | Invitrogen | L34976A |
| VPD450 |  |  | 1:400 (1,0 mg/ml stock) | BD Horizon | 562158 |
| DiO lipophilic membrane dye | | | 1:200 (5 μM stock) | Invitrogen | V22886 |
| DIL stain |  |  | 1:400 (2,5 mg/ml stock) | Invitrogen | D3911 |
| AF350 NHS Ester |  |  | 1:100 (1,0 mg/ml stock) | Invitrogen | A10168 |
| Cell trace violet | | | 1:400 (1 μM stock) | Thermo Fisher Scientific | C34557 |
| CFSE |  |  | 125nM | Invitrogen | C34554 |

**Table 2:**
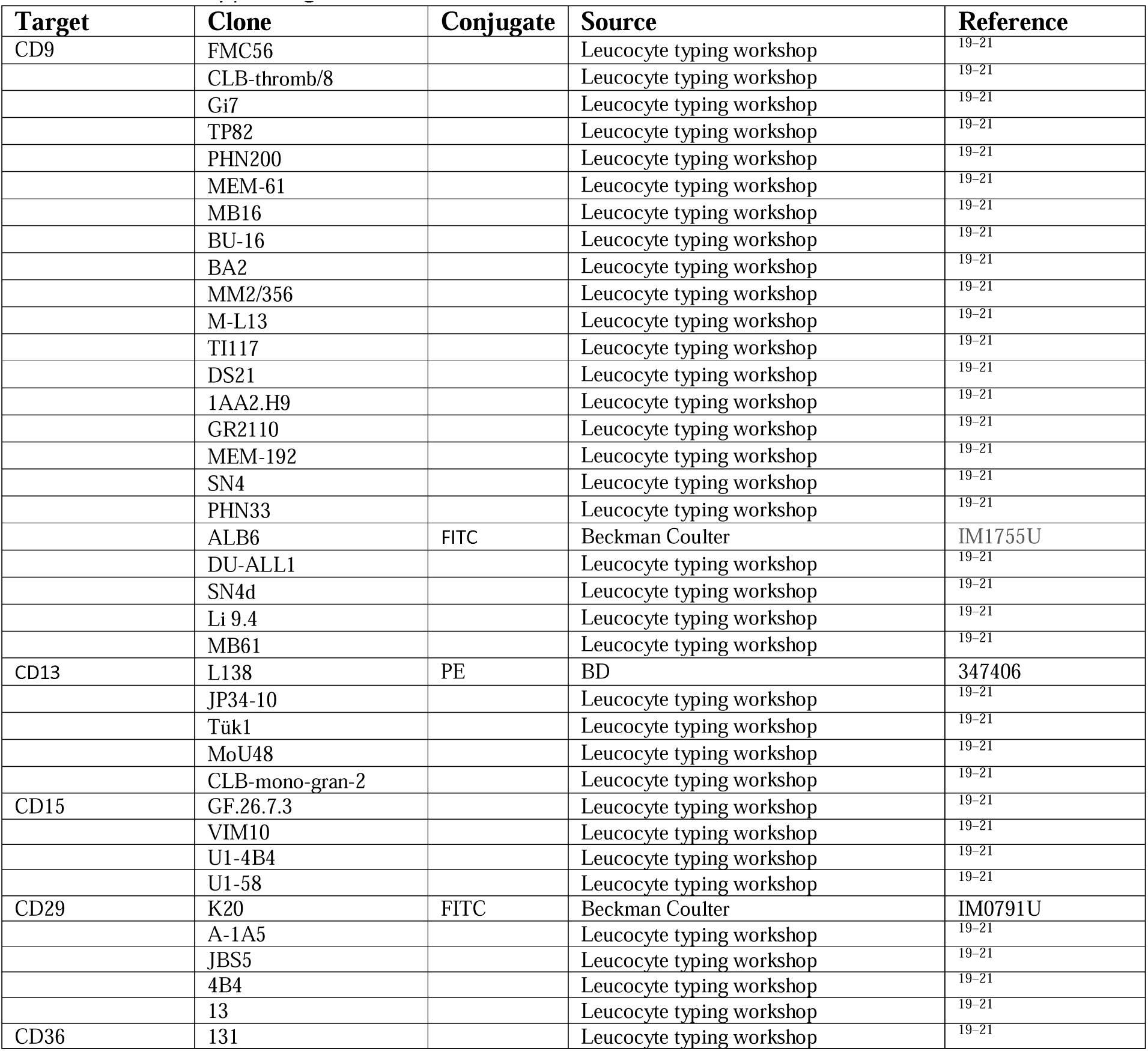
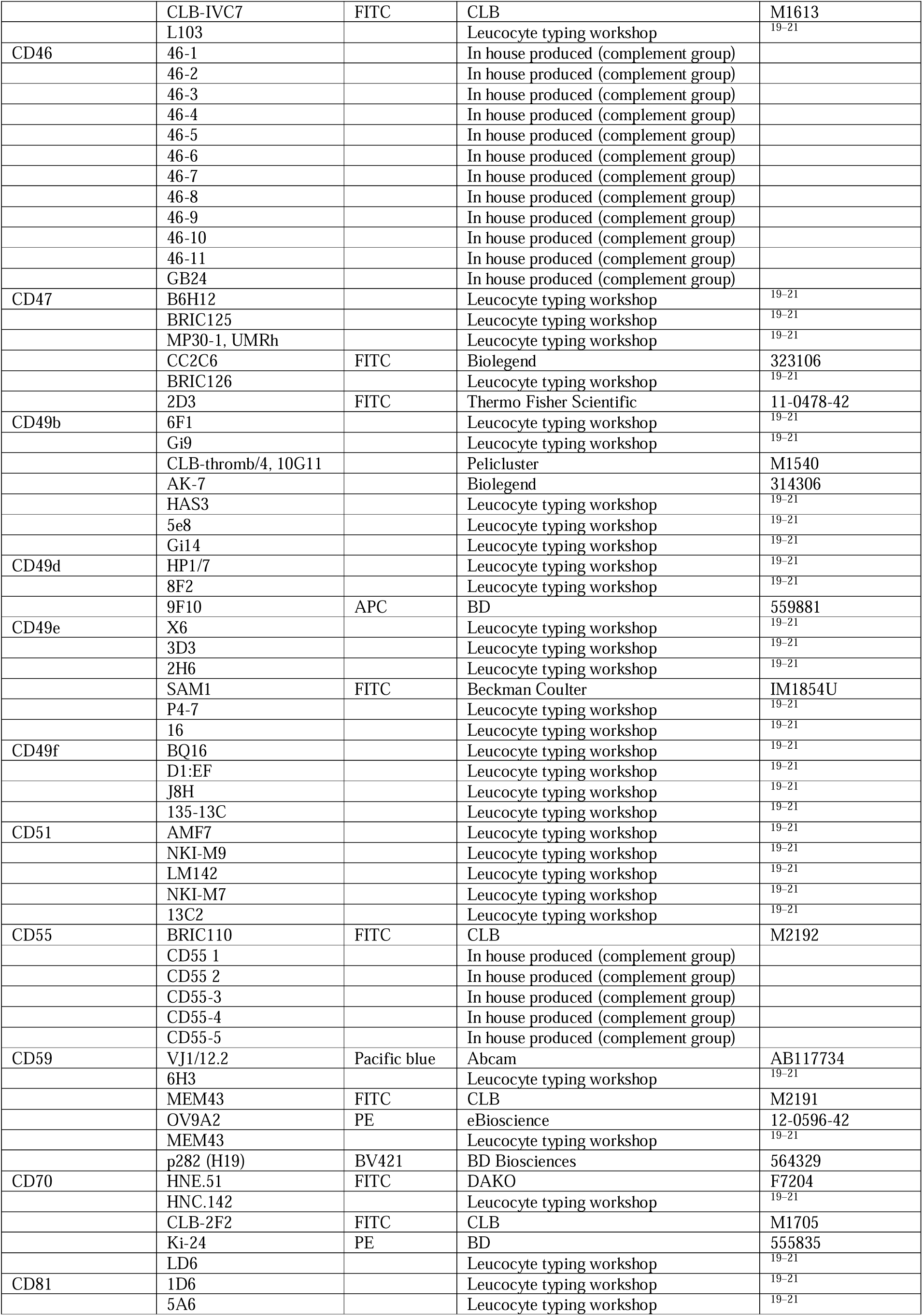
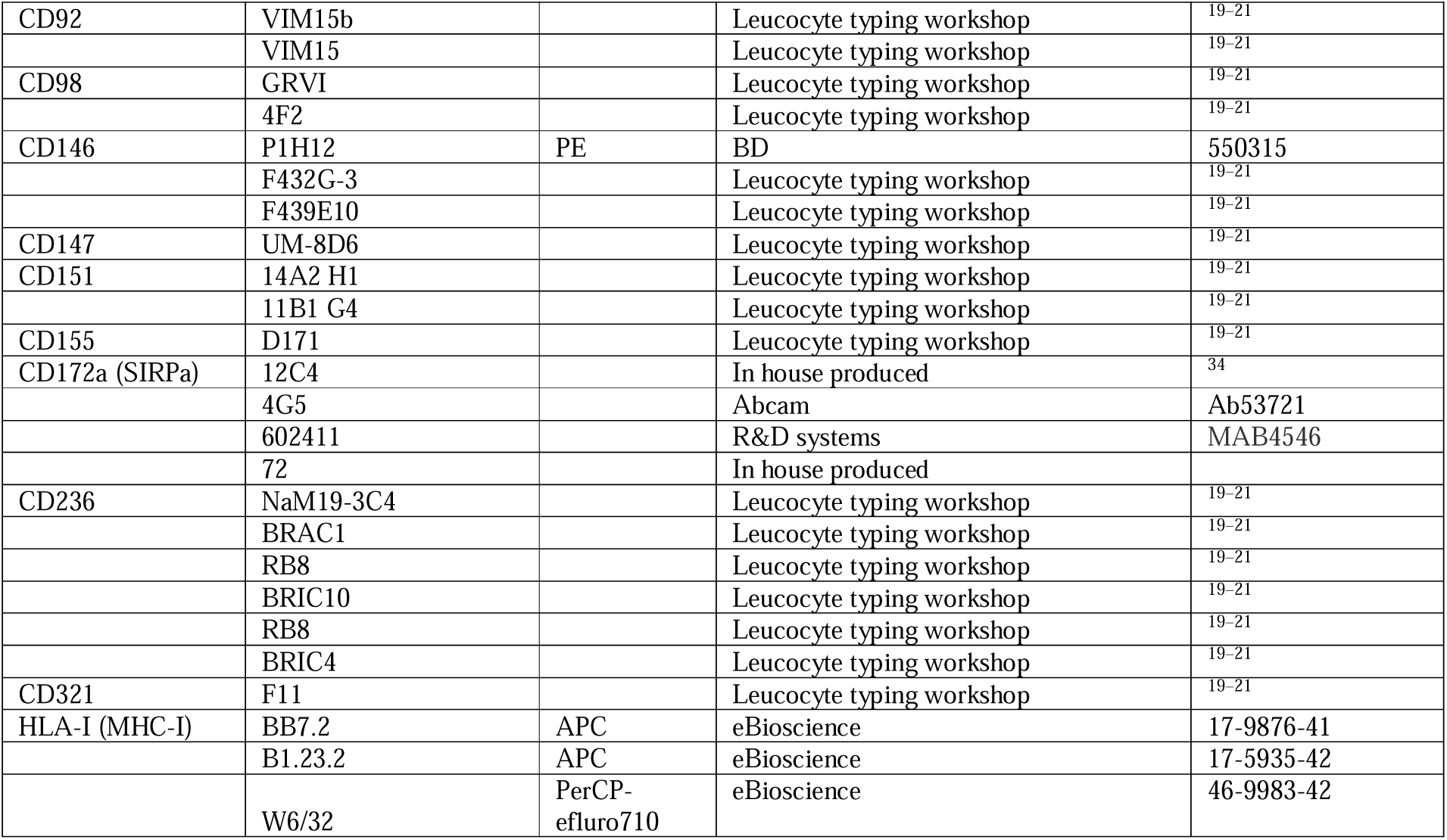
Antibody panel figure 5 and 6.

### Cell isolation and culture

#### Cell lines

HAP1 and K562 cell lines were cultured in IMDM (Gibco) and NALM6 cells in RPMI 1640 (Gibco) supplemented with 10% FCS (Serana) and 1 % antibiotics (penicillin-streptomycin, Invitrogen). HEK293T cells were cultured in DMEM supplemented with 10% FCS, 20 μg/ml gentamycin (Gibco), 1% L-glutamine (Gibco) and 0.05 mM 2-mercapto-ethanol (βME (Sigma)) and NB4 cells (kindly provided by dr. Hanke Matlung, Sanquin Research, Amsterdam, The Netherlands) were cultured in IMDM supplemented with 10% FCS, 1% L-glutamine and 1% antibiotics (penicillin-streptomycin). All cells were cultured at 37°C and 5% CO_2_.

#### Isolation of γδ T cells and NK from PBMCs

Buffy coats were acquired from Sanquin Blood Supply, Amsterdam, the Netherlands. All donors were healthy adults and had provided written informed consent in accordance with the protocol of the local institutional review board, the Medical Ethics Committee of Sanquin Blood Supply, and conforms to the principles of the Declaration of Helsinki. From these buffy coats, peripheral blood mononucleated cells (PBMCs) were isolated using Lymphoprep (Axis-Shield PoC AS, Scotland) density gradient. γδ T cells were next isolated using a direct γδ TCR targeting isolation method^16^. In short, PBMCs were incubated with a PE-conjugated mouse anti-human Vδ2 TCR (Biolegend) for 30 minutes on ice, washed with PBS/0.1%BSA and incubated with anti-mouse IgG microbeads (Miltenyi) prior to positive MACS isolation according to manufacturer’s protocol. The γδ T cells were further purified using FACS. NK cells were isolated from the PBMCs using the NK cell isolation kit from Miltenyi according to manufacturer’s protocol. The cells were negatively sorted using an LS column (Miltenyi) followed by a second LD column to remove more NK negative cells to ensure optimal purity.

#### Culture of primary cells

Purified γδ T cells were expanded for 14-days as previously described^16^. In short, the cells were cultured IMDM supplemented with 5% FCS, 5% human serum (HS, Sanquin), 1% antibiotics, 1% L-glutamine with PHA (1 µg/mL, Remel Europe), IL-2 (120 U/ml, Peprotech), IL-7 (20 ng/ml, Miltenyi) and IL-15 (20 ng/ml, Peprotech) together with feeder cells (irradiated PBMCs and Epstein-Barr virus-(EBV-) immortalized lymphoblastoid cell lines (LCLs)). After expansion, the γδ T cells were immediately used for functional assays. Isolated NK cells were cultured O/N in RPMI 1640 with 100 ng/ml IL-15 and used in functional assays the next day. Cells were cultured at 37°C and 5% CO_2_.

### Genome editing

A plasmid containing the gRNA targeting SPPL3 or UGCG (lentiCRISPR-v2-SPPL3gRNA-puro^11^) was co-transfected into HEK293T cells together with packaging plasmids psPAX2, pVSVg and pAdVAntage (Promega) using Genejammer (Agilent) for virus production. Viral supernatant was filtered and used for transduction of NALM6 and K562 cells through spinoculation together with 8 μg/ml protamine sulfate (Merck). Transduced cells were selected using puromycin (1 μg/ml).

### Functional assays

#### Coculture assays with NK cells

NALM6 or K562 target cell lines were plated two hours before coculture with NK cells in 50 μl complete RPMI (10% FCS and 1% antibiotics), at a density of 30,000 cells/well in a round-bottom 96 well plate. Next, NK cells were added to the NALM6 cells at an effector to target (E:T) ratio of 10:1, 5:1, 2.5:1, 1:1 and 0.5:1 or to K562 cells at a 5:1, 1:1 and 0:1 ratio in a total volume of 100 μl complete RPMI. After a 5-hour coculture, cells were harvested and transferred to a V-bottom plate, after which the percentage of dead target cells was determined using flow cytometry. NK cells were distinguished using anti-CD16 (Biolegend) and anti-CD56 (BD Horizon). The percentage of dead target cells observed in wells without NK cells was subtracted from the percentage of dead target cells in wells with NK cells to determine the percentage of target cell death due to coculture with NK cells.

#### Coculture assays with γδ T cells

HAP1 target cell lines were plated one day prior to the coculture with γδ T cells at a density of 15,000 cells/well in a flat-bottom 96-well plate and treated O/N or not with 10 µM pamidronate (PAM, Sigma) in complete IMDM (includes 10% FCS, 1% antibiotics). After the O/N culture, the wells were carefully washed to remove dead cells and γδ T cells were added at a 1:1, 5:1 and 10:1 ratio (E:T) in a total volume of 100 μl complete IMDM. After a five-hour coculture, the cells were harvested and transferred to V-bottom plates. Target cells that remained attached after initial harvest were detached using trypsin followed by PBS, added to the V-bottom plate and washed altogether. The percentage of dead target cells and the magnitude of γδ T cell activation were measured using flow cytometry. The γδ T cells were identified with anti-CD3 (BD Horizon) and anti-Vδ2 TCR (Biolegend), which is the same antibody used to sort the γδ T cells prior to expansion, and analyzed for CD25, CD69 and granzyme B expression. The percentage of dead target cells, gated on the γδ T cell negative fraction, observed in wells without γδ T cells was subtracted from the percentage of dead target cells in wells with γδ T cells to determine the percentage of target cell death due to coculture with γδ T cells.

#### Trogocytosis assays

Prior to the assay, 0.5·10^6^ NB4 cells/ml were differentiated with 5 μM all-*trans* retinoic acid (ATRA; Sigma-Aldrich) for seven days to promote expression of the dimer CD11b/CD18 (MAC-1) which is required for trogocytosis^17,18^. As a readout for proper maturation, expression of CD11b on stimulated NB4 cells was determined by flow cytometry. Next, NB4 cells were labeled with 2.5 μM Violet proliferation dye 450 (VPD450, BD) for 30 min at RT. The HAP1 cells were labeled with 5 μM DiO lipophilic membrane dye (Invitrogen) for 30 min at RT. After labeling and washing, HAP1 cells were incubated together with NB4 cells at 37°C in a 96 well U-bottom plate at a 5:1 E:T ratio up to four hours. Samples were fixed with stop-buffer containing 0.5% PFA and 1% BSA in PBS after one, two and four hours. The percentage of NB4 cells positive for DiO was assessed by flow cytometry.

### Flow cytometry

All antibodies and membrane dyes used are listed in *table 1 and 2*.

Prior to staining, cells were washed with PBS/0.1%BSA. Extracellular staining was performed through incubation with specific antibodies and LIVE/DEAD NEAR-IR (Invitrogen) diluted in PBS for 30 min. on ice in the dark. Next, cells were washed twice with PBS/0.1% BSA and either resuspended in PBS and directly analyzed or fixated using the BD Cytofix/Cytoperm fixation and permeabilization kit (BD Bioscience) according to the manufacturer’s protocol. For intracellular staining, cells were permeabilized and incubated with antibodies diluted in permeabilization buffer for 30 min. on ice in the dark. Subsequently, cells were washed twice and resuspended in PBS/0.1% BSA prior to analysis. Stained cells were analyzed on BD flow cytometers (LSR-II, Fortessa, FACSymphony) or sorted (ARIA-II) and analyzed using FlowJo^TM^ software version 10.9.0 (Ashland, OR: Becton, Dickinson and Company; 2023).

#### Protein panel methods, quality control and analysis

Prior to antibody staining, HAP1 WT cells were labelled with cell trace violet (Thermo Fisher Scientific), HAP1 SPPL3^-/-^ cells with AF350 succinimidyl ester (Invitrogen) and HAP1 SPPL3^-/-^B3GNT5^-/-^ with both according to manufacturer’s protocol. HAP1 B3GNT5^-/-^ cells remained unlabelled. Cells were washed thoroughly and mixed in a 1:1:1:1 ratio. Subsequently, the mixture of cells was incubated with different dilutions of 168 unique, primary antibodies which come from the Human Leukocyte Differentiation Antigen Typing Workshops, commercial suppliers or are in house produced (listed in Table 2), targeting 34 cell surface proteins for 30 min. on ice^19–21^. Cells were washed twice followed by staining with anti-mouse IgG-AF647 (Invitrogen) for 30 min. on ice in the dark. The cells were washed and resuspended in PBS/0.1% BSA for analysis.

Antibodies were excluded from analysis if there was no positive stain compared to the secondary antibody only control. Only proteins of which was established that their cell surface expression itself was not altered in one of the cell lines, as established by using saturating concentrations of at least one antibody clone per target protein. To determine whether epitope accessibility was affected by nsGSLs, the MFI for each antibody at a sub-saturating concentration was compared between the different cell lines. Epitopes were considered to be affected by the loss of SPPL3 if antibody binding to HAP1 SPPL3^-/-^ cells was different compared to WT at non-saturating antibody concentrations (MFI SPPL3^-/-^ /MFI WT <0.8 or >1.2) and by nsGSLs if (additional) knockout of B3GNT5 showed antibody binding comparable to or higher than WT levels (MFI B3GNT5^-/-^/MFI WT >0.8 or MFI SPPL3^-/-^B3GNT5^-/-^ /MFI WT >0.8).

#### Flowcytometry analysis of the CD147 targeting antibody panel

HAP1 WT cells were labelled with AF350 succinimidyl ester, HAP1 SPPL3^-/-^ cells with cell trace violet and HAP1 SPPL3^-/-^B3GNT5^-/-^ with CFSE (Invitrogen) according to manufacturer’s protocol. HAP1 B3GNT5^-/-^ cells remained unlabelled. Cells were washed twice with PBS/0.1% BSA, mixed in a 1:1:1:1 ratio and incubated with LIVE/DEAD NEAR-IR for 30 min. on ice. Cells were washed twice and incubated with 13 different anti-CD147 targeting antibodies in 50 μl PBS^22^, at concentrations of 20 μg/ml, 1 μg/ml or 0.05 μg/ml, and incubated for 30 min. on ice. Hereafter, cells were washed twice and resuspended in 35 μl goat-anti-mouse IgG (Invitrogen) in PBS for 30 min. on ice. Once more, cells were washed twice and resuspended in PBS/0.1%BSA prior to analysis using flowcytometry. Non-saturating antibody concentrations were selected for analysis. The effect of SPPL3^-/-^ on the binding of the antibodies was presented as the ratio MFI SPPL3^-/-^/MFI WT.

#### Flowcytometry using fusion proteins

Fusion proteins LIR1-Fc, NKG2D-Fc, SIRPα-Fc and Siglec7-Fc (R&D Biosystems) were reconstituted in PBS (250 μg/ml) prior to use. HAP1 cell lines were trypsinized, washed with PBS/0.1%BSA and WT cells were labelled with DIL membrane dye (Invitrogen), HAP1 SPPL3^-/-^ cells with VPD450 membrane dye (BD) and HAP1 SPPL3^-/-^B3GNT5^-/-^ with CFSE according to manufacturer’s protocol while the B3GNT5^-/-^ remained unlabelled. Next, the cells were incubated with LIVE/DEAD NEAR-IR diluted in PBS for 30 min. on ice in the dark. The cells were washed twice with PBS/0.1% BSA and mixed in a 1:1:1:1 ratio. Hereafter, cells were resuspended in 40-48 μl of a dilution series of the fusion protein solution (starting at 12 μg or 10 μg, twofold dilutions) for 60 min at RT protected from light. Cells were washed twice to remove unbound proteins and incubated with APC anti-human IgG (Biolegend) for 30 min on ice and protected from light. After incubation, cells were washed with PBS/0.1%BSA and resuspended in PBS/0.1% BSA prior to analysis.

### In silico protein height measurement

The maximal distance between amino acids of the extracellular domain of the cell surface proteins and the first amino acid after the transmembrane region was modelled using PyMOL (Schrödinger LLC, version 2.5.8) with protein structures imported from the Protein Data Bank (PDB) modeled with AlphaFold. The distance in Ångström (Å) between the alpha-carbons of all possible amino acid residue pairs and the first amino acid after the transmembrane region was calculated and the longest distance was selected from this matrix. For the integrins CD49b, CD49e and CD51 the inactive conformation was used to model maximal distance.

### Glycosylation analysis

NALM6 WT and SPPL3^-/-^ as well as K562 WT, UGCG^-/-^ and SPPL3^-/-^ were prepared for- and analyzed with porous graphitized carbon (PGC) LC-MS as described previously^11^.

### Statistical analysis

Statistical testing was done by a student’s T-test or a one-way ANOVA followed by a Tukey’s multiple comparison test. The statistical analysis was performed using GraphPad Prism version 10.0 for Mac OS (GraphPad Software, Boston, Massachusetts USA). Differences were considered significant when p≤0.05.

## Results

### SPPL3 affects anti-tumor cell cytotoxicity of **γδ** T cells

Previously, we demonstrated that the loss of SPPL3 in tumor target cells reduced effector cytokine production and tumor cell clearance by αβ T cells due to nsGSL mediated shielding of HLA-I ^11^. To assess whether SPPL3 affects the target cell recognition and anti-target cell cytotoxicity of γδ T cells, γδ T cells were expanded from PBMCs and cocultured with target HAP1 cells. Killing of HAP1 SPPL3^-/-^ cells by γδ T cells was reduced (Figure 1A, B). This SPPL3-dependent reduction in cytotoxicity was alleviated in the absence of the nsGSL producing enzyme B3GNT5 (SPPL3^-/-^ B3GNT5^-/-^) (Figure 1A, B), demonstrating that the insensitivity to γδ T cells cytotoxicity of SPPL3^-/-^ target cells involve nsGSLs. To stimulate tumor cell killing, target cells were in some cases pretreated with pamidronate (PAM)^16^. PAM pretreatment of the WT and SPPL3^-/-^ target cells made both cell types more susceptible to killing by γδ T cells, but the reduced cytotoxicity of γδ T cells against SPPL3^-/-^ compared to WT target cells remained (Figure 1A, B). A lower expression of the activation marker CD25 was observed on the γδ T cells after coculture with SPPL3^-/-^ cells compared to WT or SPPL3^-/-^B3GNT5^-/-^ cells (Suppl. Figure 1A, B). In contrast, their expression of CD69 was elevated after incubation with SPPL3^-/-^ target cells compared to the other target cells (Suppl. Figure 1A, D, E). Granzyme B levels remained unchanged (Suppl. Figure 1A, C).

**Figure 1.**
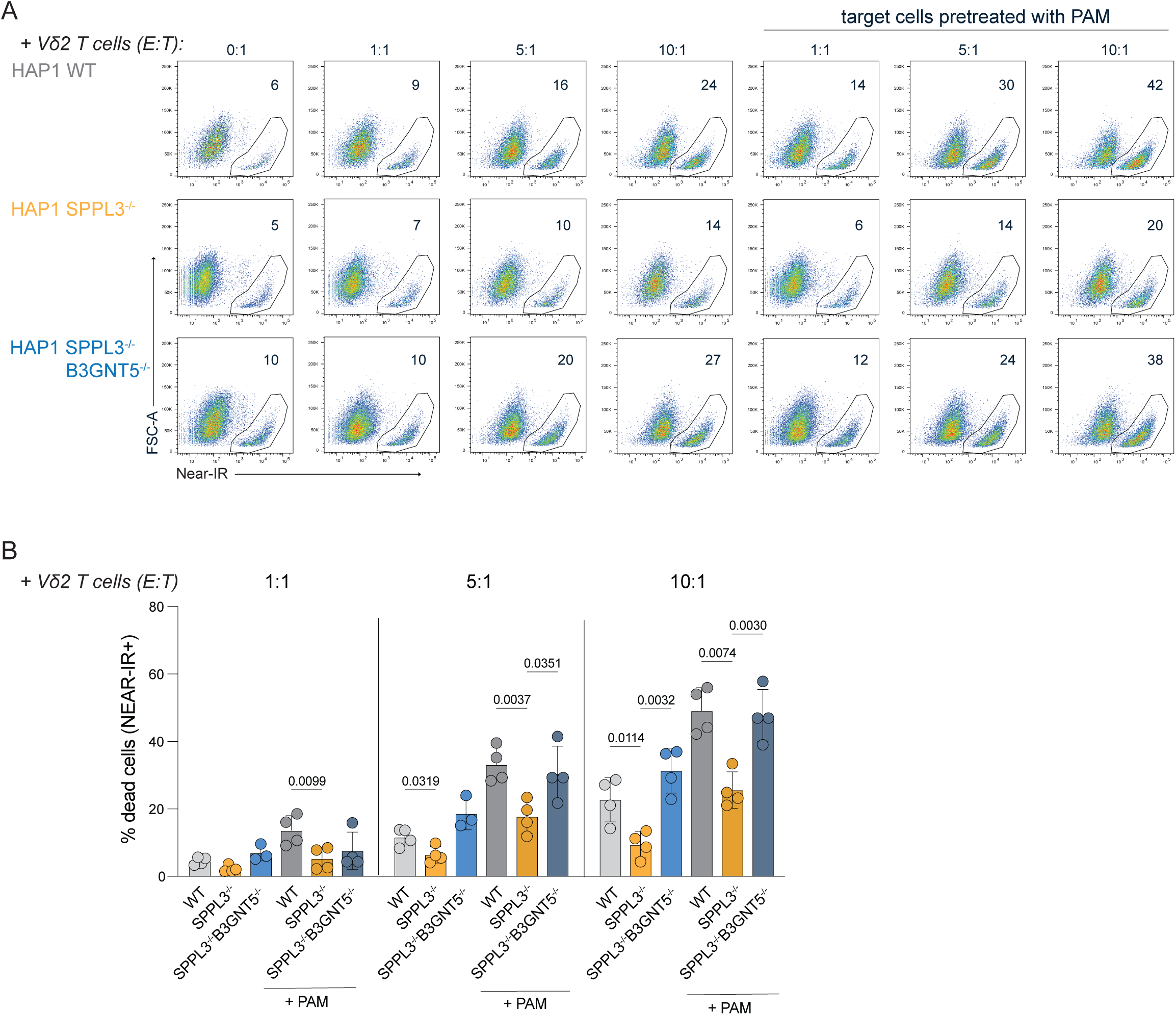
nsGSLs impair anti-tumor responses of γδ T cells. Freshly expanded γδ T cells from four donors were incubated with HAP1 cells (WT, SPPL3^-/-^ and SPPL3^-/-^B3GNT5^-/-^) for five hours in a 1:1, 5:1 and 10:1 effector to target (E:T) ratio. The target cells were either untreated or pretreated with PAM O/N prior to culture. Triplicates were used for each donor. Representative flow cytometry plots **(A)** and combined data of n=4 **(B)** showing the percentage of dead target cells (Near-IR positive) without and with addition of γδ T cells in different ratios. Each datapoint represents the average of triplicates. A one-way ANOVA was used to assess statistical significances.

Together, the data shows that the loss of SPPL3 and subsequent upregulation of nsGSLs by tumor cells protects them from elimination by γδ T cells and promotes CD69 upregulation by γδ T cells.

### SPPL3 regulates anti-tumor cell cytotoxicity of NK cells through a different pathway

To assess whether, similar to γδ T cells, NK mediated killing can be influenced by loss of SPPL3 in a nsGSL-sensitive fashion, the killing effectivity of K562 SPPL3^-/-^ and NALM6 SPPL3^-/-^ target cells by NK cells and a potential involvement of nsGSLs was investigated.

Coculture of isolated primary NK cells with WT or SPPL3^-/-^ K562 cells at different effector to target ratios demonstrated a superior killing activity against WT compared to SPPL3^-/-^ cells (Figure 2A, B), in line with the previous report. Similar results were observed for NK cocultures with WT and SPPL3^-/-^ K562 cells (Figure 2C, D). To investigate if the observed effect was due to a difference in the GSL repertoire on the NALM6 SPPL3^-/-^ cells, both WT and SPPL3^-/-^ cells were treated with the GSL synthesis inhibitor Eliglustat and stained with the antibody W6/32 targeting HLA-I which was previously shown to be inhibited by nsGSLs. Control SPPL3^-/-^ HAP1 cells recapitulated the loss of staining after Eliglustat treatment, but staining of SPPL3^-/-^ NALM6 cells remained decreased compared to WT NALM6 cells (Figure 2E)^11^. In addition, the GSL profiles of the K562 and NALM6 cells were analyzed for nsGSL expression using PGC LC-MS. K562 UGCG^-/-^ cells were taken along as a control since UGCG is a key enzyme required for GSL production^11^. Loss of SPPL3 did not promote upregulation of nsGSLs in these cells (Suppl. Figure 2).

**Figure 2.**
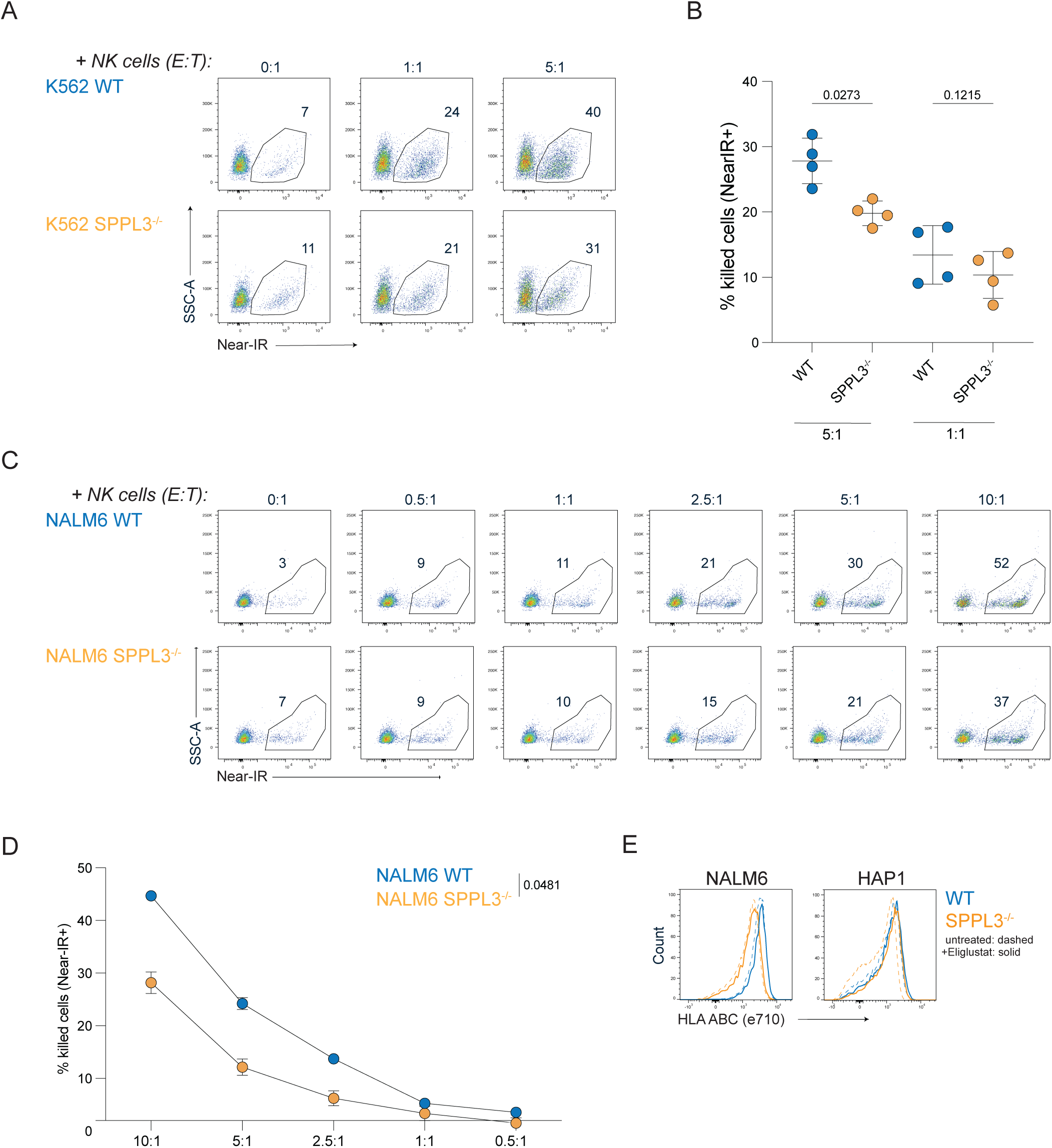
SPPL3 regulates NK cell immune responses independent of nsGSLs. NK cells were cocultured with NALM6 or K562 target cells for 5 hours in different E:T ratios. Representative flow cytometry plots **(A)** and combined data of n=2 **(B)** showing the percentage of dead (Near-IR positive) NALM6 WT or SPPL3^-/-^ cells with different E:T ratios. The datapoints represent the average of two donors for which triplicates were used. **(C)** Histograms of HAP1 and NALM6 WT (blue) and SPPL3^-/-^ (yellow) cells stained for HLA-I either untreated (dashed line) or treated with UGCG inhibitor Eliglustat (solid line) for three days. Representative flow cytometry plots **(D)** and combined data of n=4 **(E)** illustrating the proportion of dead K562 WT and SPPL3^-/-^ cells (Near-IR positive) cocultured with NK cells in a 0:1, 1:1 and 5:1 ratio. Each datapoint represents the average of triplicates. A paired student’s T-test (B) or a one-way ANOVA (E) was used to assess statistical significances.

These data showed that killing of tumor cells by NK cells is sensitive to the loss of SPPL3, however, this is probably not caused by elevated nsGSLs on the cell surface.

### Trogocytosis of SPPL3^-/-^ cells by neutrophil-like NB4 cells is diminished

Next, we investigated whether the loss of SPPL3 in target cells has an effect on neutrophil functionality, another innate immune cell type that is described to mediate anti-tumor reactivity through trogocytosis of cell membrane fragments from tumor cells ^17^. This process is initiated by the binding of the CD11b/CD18 (MAC-1) dimer on neutrophils to ICAM1 expressed by target cells. Therefore, we evaluated the effect of SPPL3 deletion on antibody binding to ICAM1 on HAP1 target cells. SPPL3^-/-^ cells showed lower cell surface staining for ICAM1 compared to WT cells, which was alleviated by the additional depletion of nsGSLs (Figure 3A). To investigate whether this difference in ICAM1 accessibility is related to affected trogocytosis, neutrophil-like NB4 cells were stimulated with all-trans retinoic acid (ATRA) for seven days to promote CD11b/CD18 expression and maturation (Suppl. Figure 3). Trogocytosis after coculture with HAP1 cells, as measured through HAP1-derived DiO-label acquisition by NB4 cells over time, showed that the efficiency of SPPL3^-/-^ HAP1 cells was lower compared to trogocytosis of WT cells (Figure 3B, C). Additional deletion of B3GNT5 improved the trogocytosis efficiency of HAP1 SPPL3^-/-^ cells. Together, these data show that SPPL3 can affect target cell trogocytosis by neutrophils, which is mediated by nsGSLs (Figure 3B, C).

**Figure 3.**
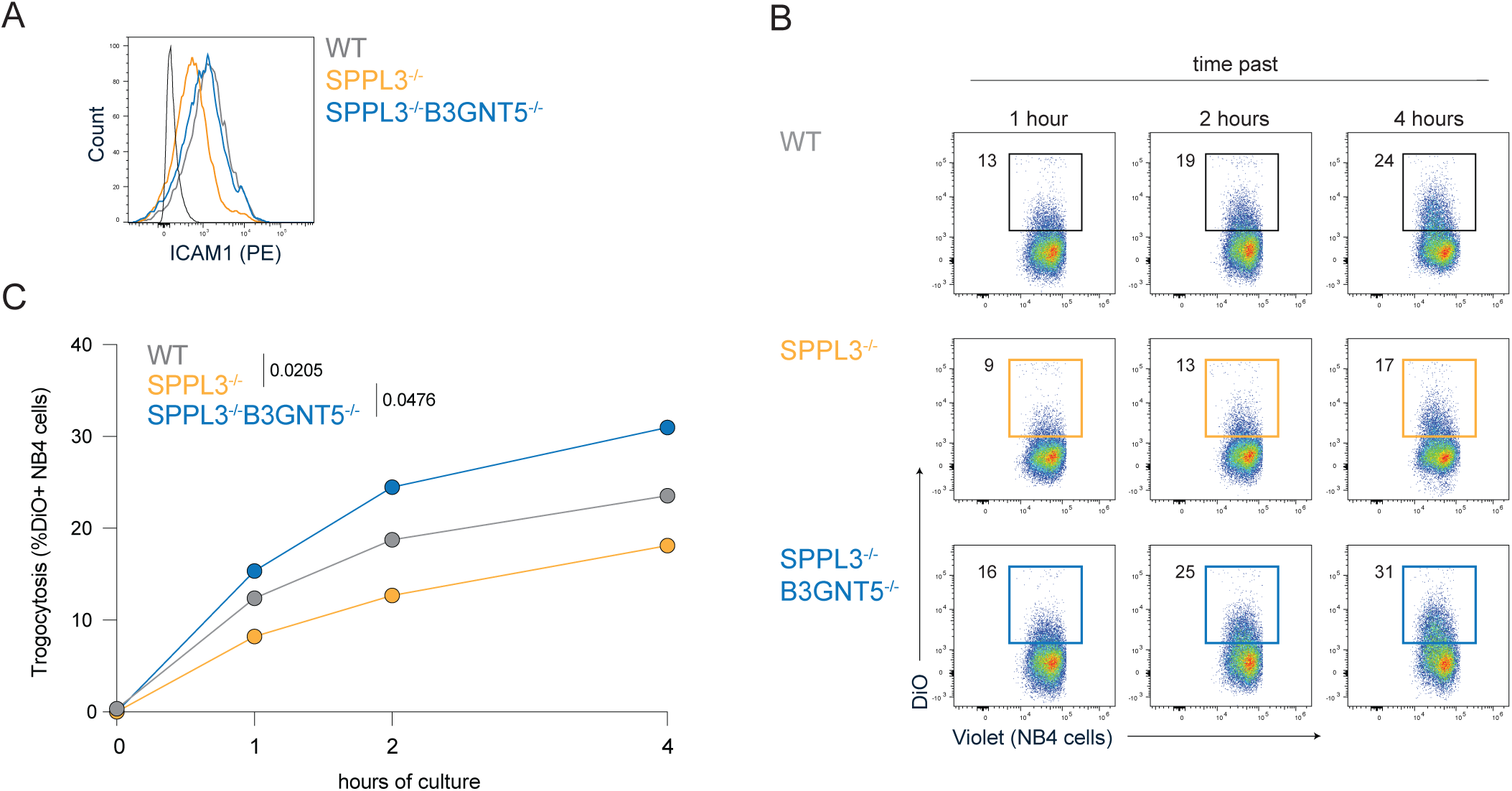
Trogocytosis of HAP1 SPPL3^-/-^ cells is diminished due to nsGSLs. (**A**) Histogram illustrating ICAM1 staining of HAP1 WT, SPPL3^-/-^ and SPPL3^-/-^B3GNT5^-/-^ cells. Trogocytosis of HAP1 cells (stained with a DiO dye) was determined by the gain of the DiO dye by NB4 cells (stained with a violet membrane dye) over a time. Representative flow cytometry plots (**B**) and combined data **(C)** where each dot represents triplicates. A paired one-way ANOVA was used to assess statistical significances.

### Loss of SPPL3 affects receptor-ligand interactions through nsGSL-dependent and independent mechanisms

Given that nsGSL-mediated inhibition of interactions is dependent on its affinity, we then investigated to which extent nsGSLs influence receptor-ligand interactions which are generally lower affinity than antibody-protein interactions. Therefore, fusion proteins of a panel of different immune receptors were used that consist of an Fc tail fused to the extracellular domain (ECD). We used fusion proteins of the phagocytosis inhibitory receptor SIRPα (ligand: CD47), expressed by myeloid cells such as neutrophils and macrophages, Siglec-7 which is a sialic acid binding inhibitory receptor mainly expressed on NK cells and to a lower extend on CD8 T cells and monocytes and the activating receptor NKG2D (ligand: MICA/B, ULBPs), expressed by NK, NKT, γδ T cells and activated macrophages^23,24^.

Previously, binding of a Leukocyte Immunoglobulin Like Receptor B1 (LIR1, LILRB1) fusion protein to HLA-I was shown to be highly impaired by nsGSLs^11^. To validate that the method used here allows for identification of differential binding in the presence of many nsGSLs, the impaired binding of LIR1-Fc to HAP1 SPPL3^-/-^ was confirmed (Figure. 4A-C).

**Figure 4:**
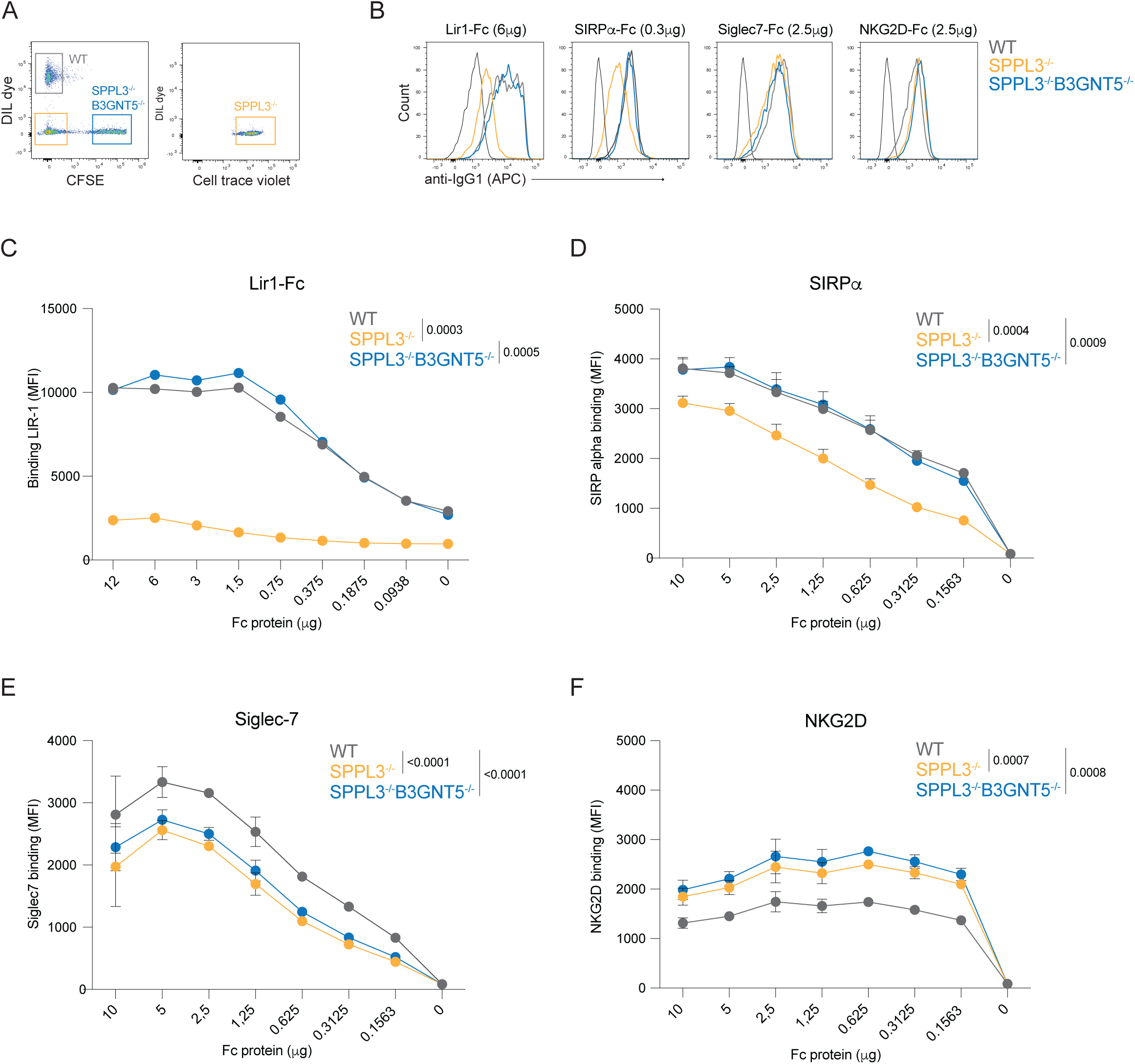
SPPL3 regulates Fc-protein binding to their ligands on HAP1 WT, SPPL3^-/-^ and SPPL3^-/-^B3GNT5^-/-^ cells. HAP1 WT (DiL dye), SPPL3^-/-^ (Celltrace violet) and SPPL3^-/-^B3GNT5^-/-^ (CFSE) were mixed **(A)** and incubated with different concentrations of LIR1-Fc, SIRPα-Fc, Siglec-7-Fc or NKG2D-Fc. Representative flow cytometry plots **(B)** and summarizing graphs for the binding of LIR-1-Fc **(C)**, SIRPa-Fc **(D)**, Siglec-7-Fc **(E)** and NKG2D-Fc **(F)**. Each dot represents the average of triplicates. A paired one-way ANOVA was used to assess statistical significances.

The interaction of the SIRPα-Fc with CD47 was impaired after loss of SPPL3. This could be restored by deletion of B3GNT5, indicating that nsGSLs impaired binding of SIRPα (Figure 4B, D). The binding of Siglec-7 to SPPL3^-/-^ cells was reduced independently of nsGSLs produced by B3GNT5 (Figure 4B, E). Likewise, the interaction between NKG2D and probable binding partners MICA/B and/or ULPBs is improved with the loss of SPPL3, independent of B3GNT5 produced nsGSLs (Figure 4B, F).

Altogether, multiple receptor-ligand interactions are modulated depending on SPPL3 expression through nsGSL-dependent and independent mechanisms.

### The accessibility of several membrane proteins is affected by nsGSLs

We previously demonstrated that enhanced nsGSLs in SPPL3^-/-^ cells affect the accessibility of several epitopes on HLA-I^11^. To assess whether the effect of SPPL3 and nsGSL expression in target cells on NK cells, γδ T cells and neutrophils is driven by the cumulative effect of different affected cell surface proteins, we assessed the effect of SPPL3 and nsGSL expression on 34 different cell surface proteins using a panel of 168 antibodies. The various HAP1 cell lines were barcoded with different membrane dyes and mixed prior to antibody staining to allow for most optimal comparison (Figure 5A). Out of the 34 proteins assessed with this panel, the antibody staining of 25 proteins passed quality control (requirements are listed in the methods, listed in Table 2). Per antibody, the ratio of the mean fluorescence intensity (MFI) between each KO cell line and the WT cell line was determined. Ratios between 0.8 and 1.25 were considered similar to WT (examples in Figure 5B left panel) and values below or above this cutoff were considered different (examples in Figure 5B right panel).

**Figure 5.**
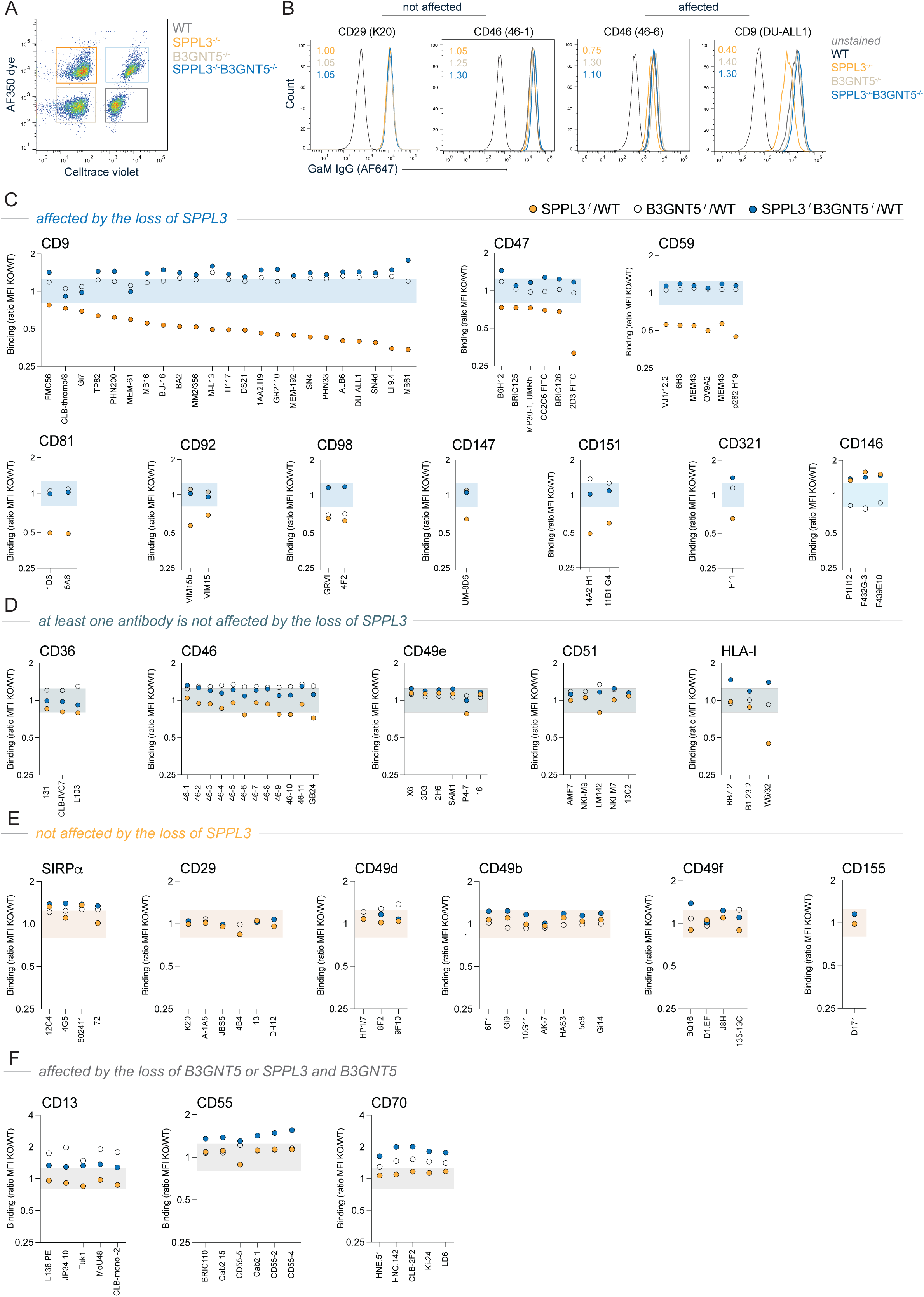
Antibody binding to proteins on HAP1 SPPL3^-/-^ and HAP1 SPPL3^-/-^B3GNT5^-/-^ relative to HAP1 WT cells. HAP1 WT (Celltrace violet), SPPL3^-/-^ (AF350 dye) and SPPL3^-/-^B3GNT5^-/-^ (Celltrace violet and AF350 dye) **(A)** were mixed and incubated with 168 unique, primary antibodies targeting 34 cell surface proteins. Antibodies targeting a total of 25 proteins passed quality control. **(B)** A proportion of proteins were targeted with antibodies that were all not affected to bind to their epitopes with SPPL3^-/-^ while some proteins were partially (at least one of the tested antibodies) or completely affected (all tested antibodies), which could be alleviated with additional B3GNT5^-/-^. The ratio of the MFI of each antibody for the SPPL3^-/-^, B3GNT5^-/-^ and SPPL3^-/-^B3GNT5^-/-^ compared to the WT cells was calculated (example in plots in B). The proteins were grouped based on the effect of the loss of SPPL3 or B3GNT5; proteins with epitopes affected by the loss of SPPL3 **(C)**, proteins of which at least one antibody is not affected by the loss of SPPL3 **(D)**, proteins with epitopes that are not affected by the loss of SPPL3 **(E)**, proteins with epitopes affected by the loss of B3GNT5 or SPPL3 and B3GNT5 **(F)**. The colored bar represents the area that is considered similar to WT (cutoff for difference: >0.8 and <1.2).

Based on this analysis, we could discriminate between proteins that were affected, not affected or at least in part not affected by loss of SPPL3 (Figure 5B, C-F). In contrast to the other proteins, accessibility of CD146 by its specific antibodies was increased in the absence of SPPL3 (Figure 5C). The group of proteins for which all antibodies tested were negatively affected by SPPL3 KO consisted of CD9, CD47, CD59, CD81, CD92, CD98, CD147, CD151 and CD321 (Figure 5C). SPPL3 effects could be reversed by additional depletion of nsGSLs, demonstrating that SPPL3 loss affected these proteins through modulation of nsGSLs.

At least one, but not all antibodies targeting the proteins CD36, CD46, CD49e and CD51 were not affected in the HAP1 SPPL3^-/-^ cells, similar to antibodies targeting HLA-I as described (Figure 5B, D)^11^. Antibody binding to SIRPα, CD29, CD49b, CD49d, CD49f and CD155 was generally equal between the tested cell lines (Figure 5B, E). A small subgroup of proteins was specifically affected by the loss of B3GNT5, but not SPPL3 (Figure 5F).

### nsGSLs do not limit accessibility to proteins with long extracellular domains

We hypothesized that antibody epitopes more distal from the cell membrane may be not subjected to shielding by nsGSLs. However, for most of the antibodies used, the exact epitopes have not been properly mapped. Therefore, we investigated whether the size of the ECD correlated with the nsGSL-dependent SPPL3 sensitivity of those proteins.

For 15 proteins the structure was modeled in AlphaFold. The distance between the alpha carbons of each amino acid within the ECD and the alpha carbon of the ECD was determined and the longest distance (Å) was used as the size of the ECD (Suppl. Figure 4, Table 3). For example, the largest ECD of CD9 has a diameter of 40.5Å (Figure 6A, Suppl. Figure 4, Table 3).

**Figure 6:**
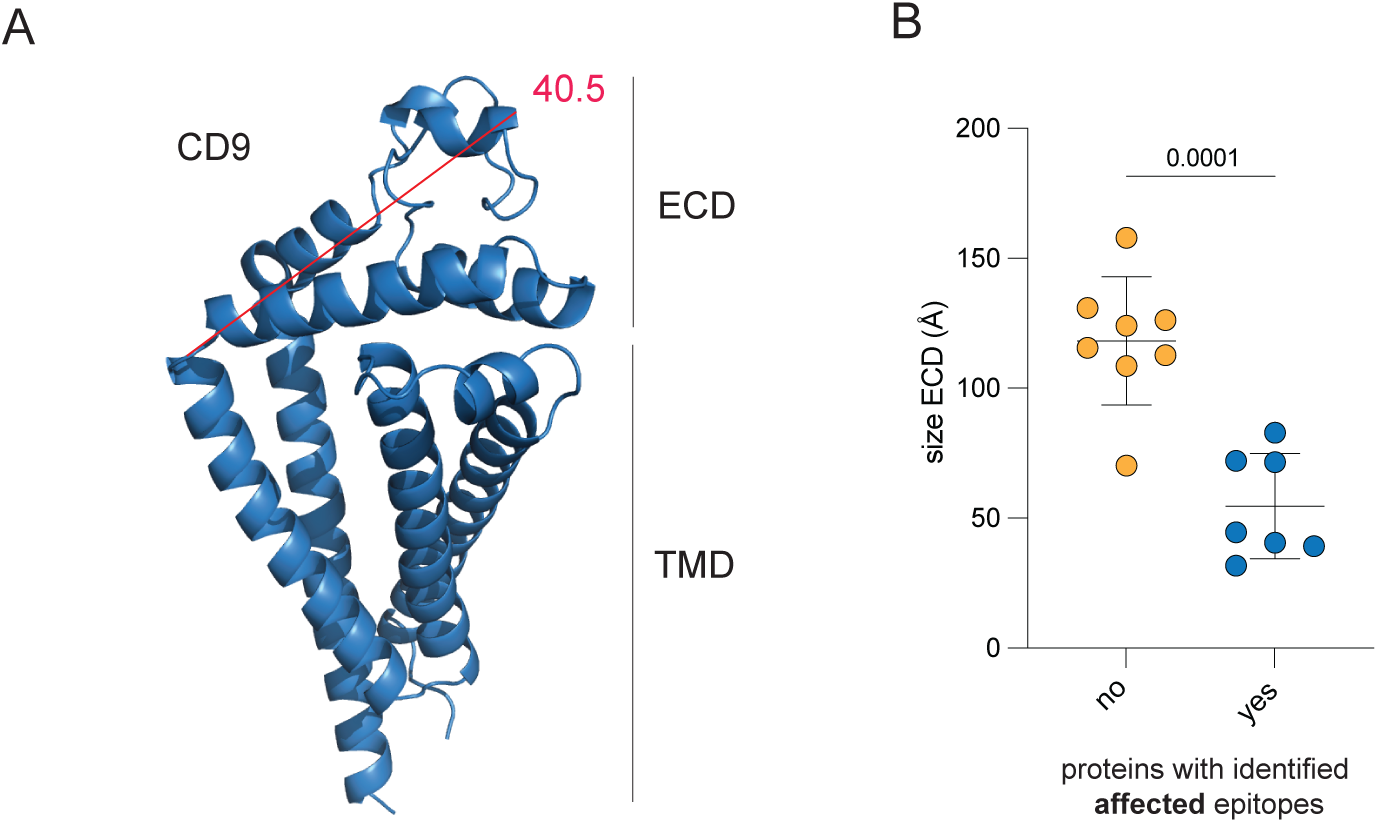
The protrusion of proteins affected by nsGSLs versus unaffected proteins. Available protein structures were obtained from the Alphafold database for the proteins that contained epitopes that were affected or not affected by nsGSLs. The distance between the alpha carbon of the first amino acid of the extracellular domain to all other amino acids was determined in Armstrong (Å) using PyMOL and the longest theoretical distance was selected. For example, **(A)** the maximum distance of CD9 is 40.5Å. **(B)** The maximum protruding distance (size) of proteins with versus the proteins without identified affected epitopes.

**Table 3.** List of affected proteins and proteins with at least one non-affected epitope. ECD: Extracellular domain.

| Affected proteins | Size ECD (Å) | Proteins with at least one non-affected epitope | Size ECD (Å) |
| --- | --- | --- | --- |
| CD9 | 40.5 | CD46 | 112.8 |
| CD47 | 44.6 | CD49b | 157.9 |
| CD59 | 31.7 | CD49d | 108.5 |
| CD81 | 39.2 | CD49e | 126.3 |
| CD98 | 72.0 | CD51 | 130.9 |
| CD147 | 71.5 | CD155 | 115.6 |
| CD321 | 82.9 | SIRPα | 124.1 |
|  |  | HLA-I | 70.2 |

The ECD size of cell surface proteins that contain domains that are not affected by nsGSLs were significantly larger (μ=126±22Å) than proteins for which we did not identify such domains (μ=55±20Å) (Figure 6B). These data indicate that the accessibility of cell membrane proteins that extend less from the plasma membrane is more prone to be affected by loss of SPPL3 compared to proteins that extend further from the plasma membrane.

### Severity of nsGSL effect is dependent on affinity of a protein interaction

We then wondered whether within the group of nsGSL affected interactions, affinity or further distance measures of such protein-protein interaction may play a role in the degree by which nsGSLs diminish binding. To test these hypotheses, we used CD147 as model antigen, since the previously tested antibody was affected by nsGSLs (Figure 5), since it has a significant ECD size of 71.5Å (table 3, Suppl. Figure 4) and since we had access to 13 different CD147-specific antibodies, each with well-defined epitope positions and binding affinities^22^. We then evaluated the loss of binding of these antibodies by nsGSLs by staining HAP1 cells with the different nsGSL profiles (WT, SPPL3^-/-^, B3GNT5^-/-^ and SPPL3^-/-^B3GNT5^-/-^). In comparison to WT cells, all 13 antibodies showed impaired binding to SPPL3^-/-^ cells (Figure 7A, Suppl. Figure 5). The impaired binding of these antibodies was alleviated by additional deletion of B3GNT5, validating that nsGSLs were the cause of reduced binding to CD147 (Figure 7A). The respective antibody binding affinity correlated strongly with the degree of compromised binding (Figure 7B).

**Figure 7:**
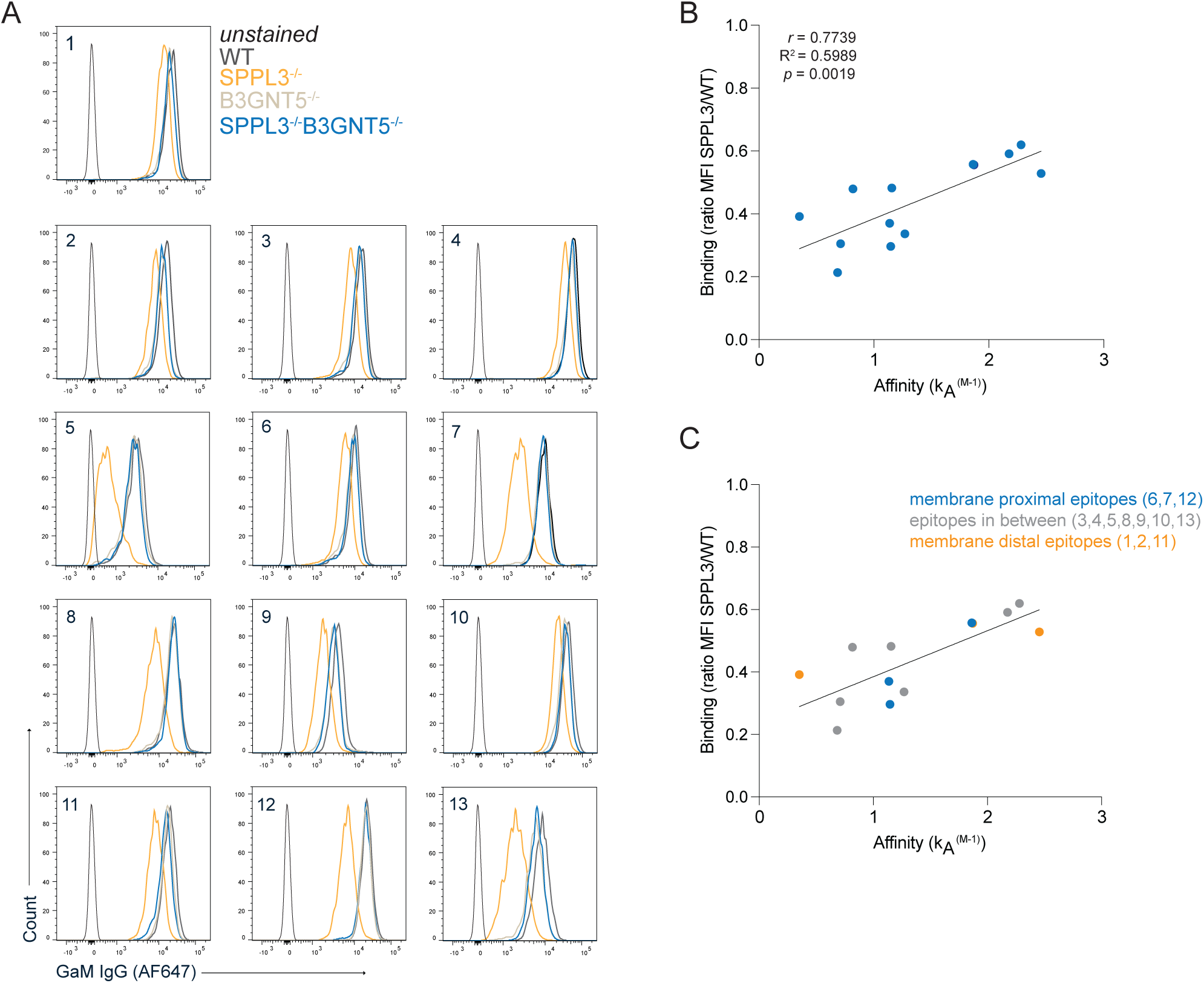
The binding of multiple anti-CD147 antibodies with different affinities to HAP1 WT, HAP1 SPPL3^-/-^, HAP1 B3GNT5^-/-^ and HAP1 SPPL3^-/-^/B3GNT5^-/-^ cells. Thirteen different antibodies targeting CD147 were incubated with ‘barcoded’ and mixed HAP1 WT (AF350 dye), SPPL3^-/-^ (Celltrace violet), B3GNT5^-/-^ (unstained) and SPPL3^-/-^/B3GNT5^-/-^ (CFSE) cells. **(A)** Representative flow cytometry plots for each antibody. **(B)** The correlation plot of the degree of SPPL3-sensitivity of binding for each antibody (SPPL3^-/-^ MFI/WT MFI) versus the affinity of the antibodies for their epitope. **(C)** Categorization of the epitopes targeted by the antibodies based on the position of the epitopes relative to the cell membrane.

Because ECD size relates to accessibility of an epitope (Figure 6), we assessed whether the position of the epitopes targeted by the CD147 antibodies in relation to the cell membrane would also play a role in the diminished binding. In their study, Koch and colleagues used a cross-blocking analysis to predict the positions of the epitopes for each of the CD147 targeting antibodies in relation to each other and the cell membrane ^22^. Antibodies six, seven and twelve target membrane proximal epitopes, one, two and eleven target membrane distal epitopes and the others target epitopes in-between. However, within this group of nsGSL-affected antibodies, their epitope positions were not related to the severity of diminished binding (Figure 7C). Together, these data suggest that the degree by which protein interactions with receptor domains are shielded by nsGSLs are additionally dictated by affinity of the interaction.

## Discussion

We showed that tumor cells that lack SPPL3 possess the ability to evade innate immune responses mediated by NK cells, γδ T cells and neutrophils. The diminished responses of the latter two are driven by increased nsGSL levels. For both nsGSL-dependent and -independent processes, the net cellular effect is likely caused by modulation of multiple receptor-ligand interactions in the absence of SPPL3. With a representative panel we showed that several membrane proteins are affected by the loss of SPPL3 in a nsGSL-sensitive fashion. Moreover, we established that the degree of nsGSL-dependent, SPPL3-sensitivity of these proteins is correlated to the size of surface receptors to the plasma membrane and the affinity between protein-protein interactions. Therefore, expression of SPPL3 by tumor cells influences crosstalk between immune cells through a multitude of receptor-ligand interactions thereby driving escape not only from adaptive but also from innate immunity.

We showed that K562 and NALM6 SPPL3^-/-^ cells were more resistant to NK cell mediated killing without involvement of nsGSLs. The target cell lines utilized in this study did not show upregulation of nsGSLs with the loss of SPPL3. Therefore, it is not concluded that nsGSLs cannot influence NK cell mediated killing, which should be tested with target cell lines in which nsGSL expression is established.

In line with our data, Dufva and colleagues identified SPPL3 as a gene that facilitates susceptibility to NK cell mediated killing^25^. Moreover, Heard and colleagues showed that binding of anti-CD19 antibodies to NALM6 SPPL3^-/-^ was impaired and, in addition, that the loss of SPPL3 in NALM6 cells increases their resistance towards CD19-targeting CAR T cells^12^. Absence of SPPL3 results in increased and altered *N*-glycosylation of CD19 which disrupts the CAR binding epitope. Both antibody binding and CAR T cell killing could be restored by using kifunensine, a potent inhibitor of mannosidase I, indicating an important role of *N*-glycosylation in the recognition of target proteins in NALM6 cells. Altered *N*-glycosylation of other ligands on tumor cells, may likewise be important for recognition by NK cells and may explain the lower NK cell mediated cell death of SPPL3^-/-^ compared to WT NALM6 cells presented in this work. Paradoxically, binding of NK cell activating receptor NKG2D to HAP1 cells improved with the loss of SPPL3 through a mechanism independent of nsGSLs. It is possible that other important ligand-receptor interactions are impeded which may favor tumor survival by sustaining the inactivation of NK cells in the functional assays. The diminished NKG2D-Fc binding on HAP1 cells, may have involved shedding of its ligands. Tumors use shedding of the ectodomains of NKG2D ligands MICA/B and ULPBs as an effective mechanism to escape from NK cell recognition^26,27^. Shedding of these proteins involves activity of metalloproteases such as a disintegrin and metalloproteinase 10 (ADAM10)^28^. SPPL3 has also been shown to act as a selective sheddase that catalyzes the release of the ectodomain of substrates^29–31^. In addition, SPPL3 can mediate activation of metalloproteinase ADAM10^32^. It is unknown if MIC proteins or ULBPs are direct substrates for SPPL3, though it could explain the increased binding of NKG2D when loss of SPPL3 would impair shedding and resulting in increased protein availability on the cell membrane.

In addition to regulation of innate immunity through *N*-glycosylation, regulation of nsGSL levels by SPPL3 greatly impacts the effective killing of tumor cells by γδ T cells and trogocytosis by neutrophil-like NB4 cells, underscoring the versatile mechanisms by which SPPL3 supports innate immune escape. In line with these results, Rigau and team identified SPPL3 in a genome-wide knockdown screen as one of the key factors involved in Vδ2 TCR tetramer binding^33^. Here, we identify the involvement of SPPL3 and downstream nsGSLs in the actual killing of tumor cells by γδ T cells, since additional knockout of B3GNT5 reestablishes killing. The relation between SPPL3 and γδ TCR engagement is yet to be elucidated, although it is not unthinkable that nsGSLs shield butyrophilins comparable to HLA-I and that this interaction contributes to the sum of all interactions.

Our previous work illustrated different degrees of impaired binding due to elevated nsGSLs for different antibodies targeting HLA^11^. We suggested that this was related to the position of the antibodies, since antibodies targeting the α3 domain (membrane proximal) were more impaired than antibodies binding epitopes on the α1 and α2 (peptide binding groove, membrane distal). In this study, we confirmed that position of the epitope plays a role by using a large set of antibodies. Comparison of proteins harboring epitopes that were affected by elevated nsGSL levels or not showed that proteins with relatively small ECD were prone to be affected over proteins with a larger ECD. With the exception of HLA-I, the exact amino acids involved in binding of the antibodies used here are unknown. Future epitope mapping and affinity determination of these antibodies may allow generation of a more complete analyses of the nsGSL-mediated shielding phenomenon.

In addition, we utilized 13 different antibodies targeting CD147 for which we also had affinity information. The data generated using these antibodies showed that within affected regions of the protein, the affinity is the main determinant for the degree of shielding. Still, precise affinity measurements of the previously used HLA-I specific antibodies or precise epitope identification of the CD147 specific antibodies could provide an even better understanding on the role of affinity and the distance of these epitopes to the cell membrane in protein-protein (receptor-ligand) interactions in the presence of elevated nsGSL levels.

Alternatively, one could design an artificial scaffold protein on which at different distances from the membrane the same epitope is engineered. Different affinity-versions of antibodies against such epitope could more definitively untangle those effects, although additional mechanisms may further contribute to nsGSL shielding. One such parameter could the outer charge of the cell surface receptor given the negative sialic acids are a requirement for nsGSL mediated shielding^11^.

The work demonstrated here and by others indicate that SPPL3 is an essential regulator of membrane receptors functions through regulation of nsGSL synthesis and *N*-glycosylation. While only a fraction of membrane proteins was examined in this study, the selection process was random, suggesting it could be representative for the majority of membrane proteins. Cell-cell interactions encompass a multitude of protein-protein interactions and the induction of multiple signaling pathways, culminating in cellular responses that hinge on the collective impact of these signals. It is not possible to assess the contribution of individual protein interactions to the described cellular immune responses, yet, the the cumulative interactome changes after loss of SPPL3 expression causes reduced reactivity by NK cells, γδ T cells and neutrophils. Though, targeting SPPL3 in therapies presents challenges due to its broad inhibitory function on glycosylation of multiple entities. Nevertheless, part of the regulation of SPPL3 goes though GSLs which could be a targeted via more specific inhibitors. Enhancing the effectiveness of both cell- and antibody-based therapies against nsGSL-expressing tumors could be achieved by specific inhibitors of GSL synthesis such as Eliglustat or Miglustat (UGCG inhibitors) which are currently safely applied to patients with lysosomal storage disorders^11^.

## Acknowledgements

We would like to thank the Sanquin Research core facility for their help, Sanquin Diagnostics and Dr. Richard Pouw (complement group, Sanquin Research) for providing us with the CD46 and CD55 antibodies and prof. Dr. Ellen van der Schoot (Sanquin Research) for sharing the Leucocyte typing workshop antibodies with us. This research was supported by grants from the Dutch Research Council (Grant NWO-VIDI 91719369 to R.M.S), KWF Alpe d’HuZes (BMA 2015–7982 to R.M.S.) and the Landsteiner Foundation for Blood Transfusion Research (LSBR Fellowship 1842F to R.M.S).

## Figure legends

**Supplemental figure 1. Coculture of NK and γδ T cells with WT or SPPL3^-/-^ tumor cells.**

**(A)** Histograms showing expression of CD25, granzyme B and CD69 by the γδ T cells after coculture with the HAP1 target cells. The MFI of CD25, granzyme B and CD69 positive γδ T cells **(B and C)** and percentage of CD69 positive cells **(E)** after coculture with HAP1 WT, SPPL3^-/-^ or SPPL3^-/-^ B3GNT5^-/-^ cells. Combined data of n=4, each datapoint represents the average of triplicates. A one-way ANOVA was used to assess statistical significances.

**Supplemental figure 2. Chromatograms of PGC LC-MS on total GSL glycans**

Extracted ion chromatograms of total GSL glycan content released from **(A)** NALM6 WT and SPPL3^-/-^ cells as well as **(B)** K562 WT, UGCG^-/-^ and SPPL3^-/-^ cells using PGC LC-MS.

**Supplemental figure 3. CD11b expression by maturated NB4 cells.**

Flow cytometry plot demonstrating the CD11b staining of NB4 cells after seven days of stimulation with ATRA.

**Supplemental figure 4. Protein structures of proteins with and without epitopes identified to be affected by nsGSLs.**

Available protein structures were obtained from the Alphafold database for the proteins that contained epitopes that were affected (blue), for which at least one antibody was affected (purple) or not affected (yellow) by nsGSLs. The distance between the alpha carbon of the first amino acid of the extracellular domain to all other amino acids were determined in Armstrong (Å) using PyMOL and the longest theoretical distance is shown (red line). ECD: extracellular domain. TMD: transmembrane domain.

**Supplemental figure 5. Gating strategy of mixed HAP1 WT, SPPL3^-/-^ and SPPL3^-/-^B3GNT5^-/-^ cells incubated with CD147 targeting antibodies.**

