## Supplementary figures and images for "Tumor-expressed SPPL3 supports innate anti-tumor immune responses"

### Supplementary figure 1

Supplemental figure 5

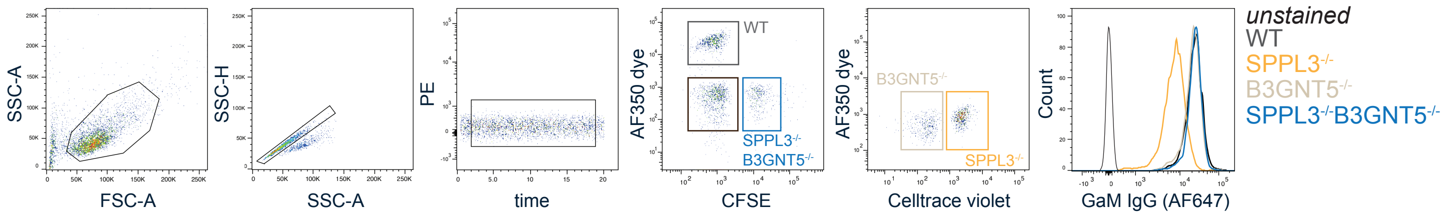

### Supplementary figure 2

Supplemental figure 3

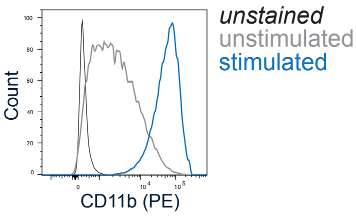

### Supplementary figure 3

Supplemental figure 4

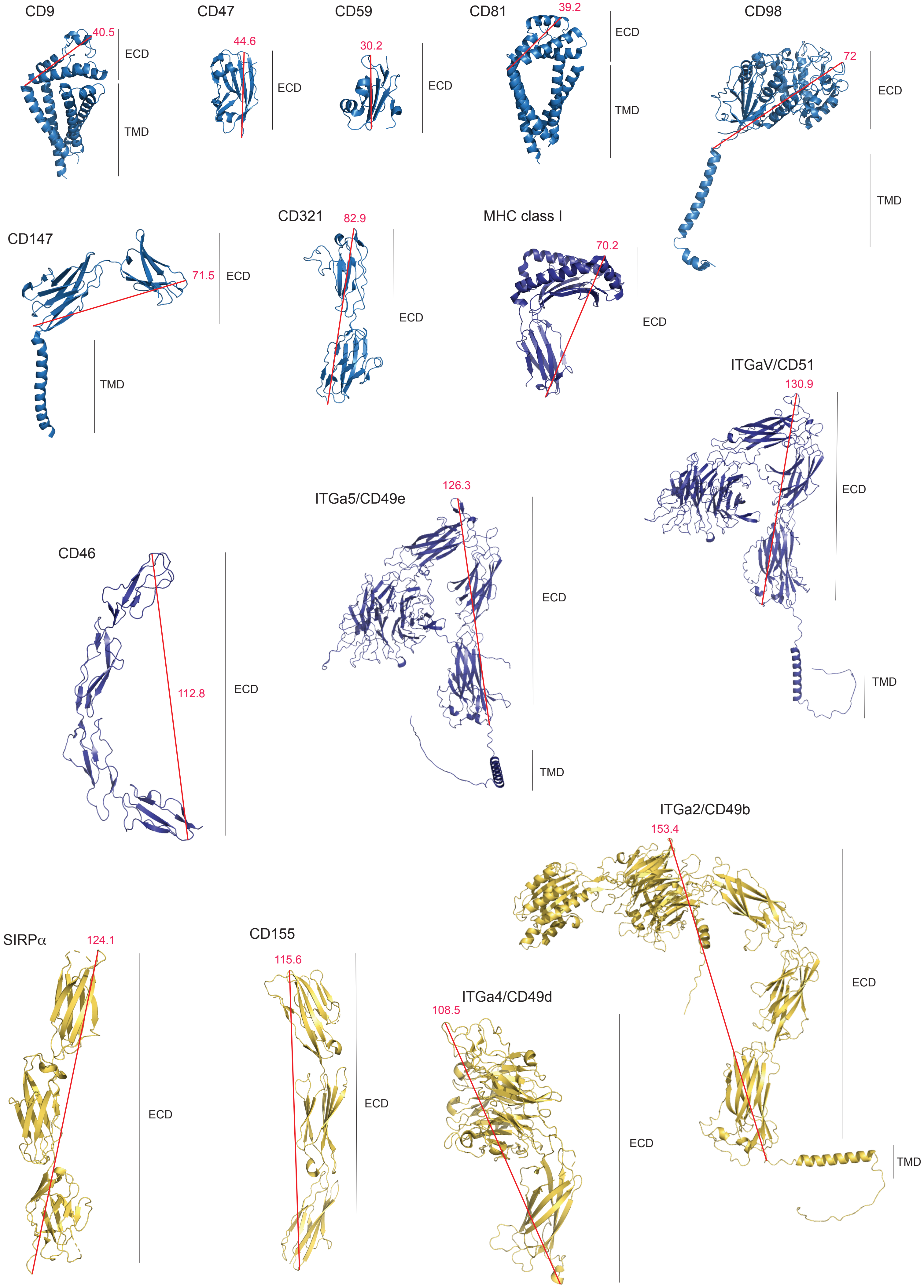

### Supplementary figure 4

Supplemental figure 2

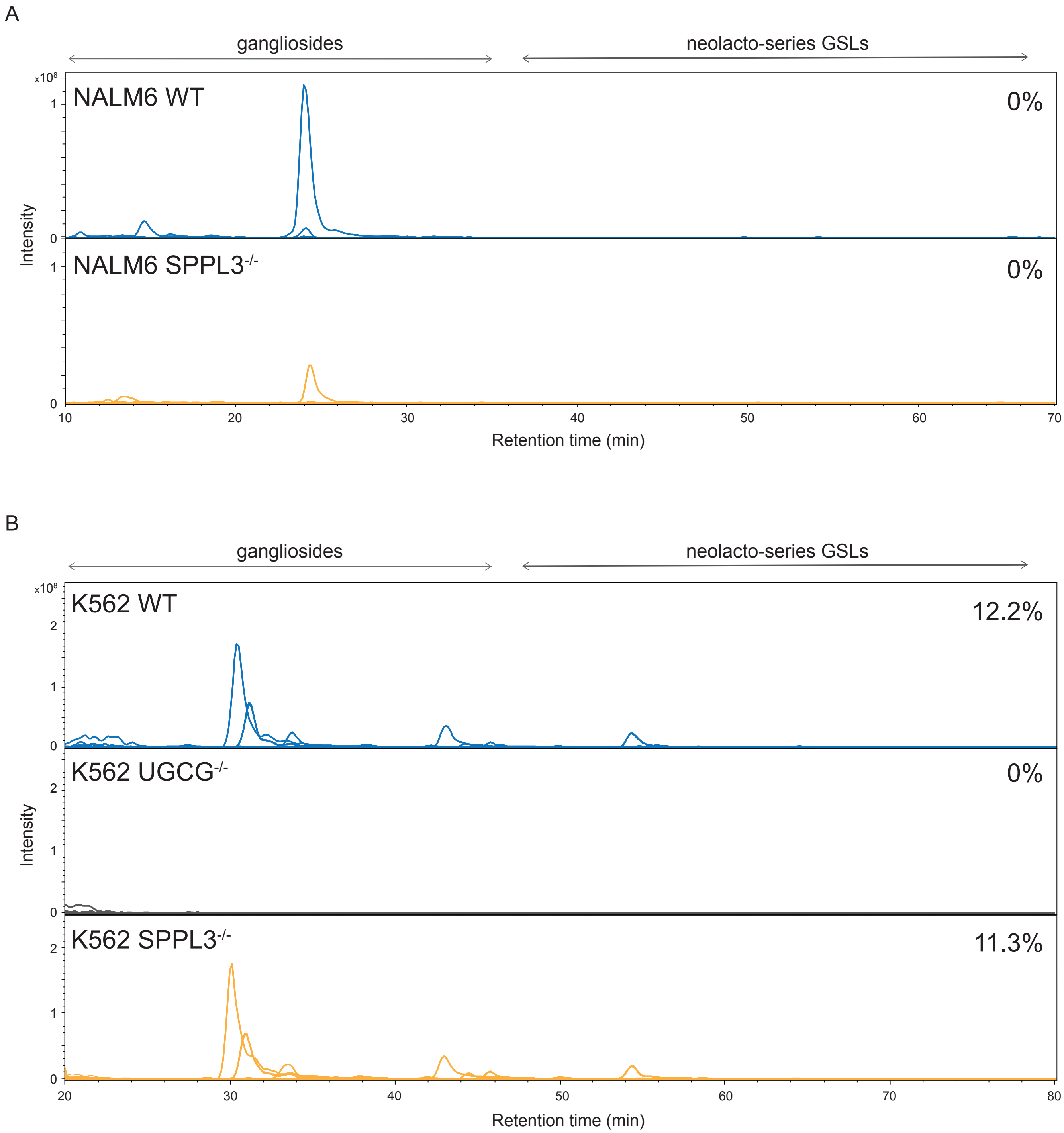

### Supplementary figure 5

Supplemental figure 1

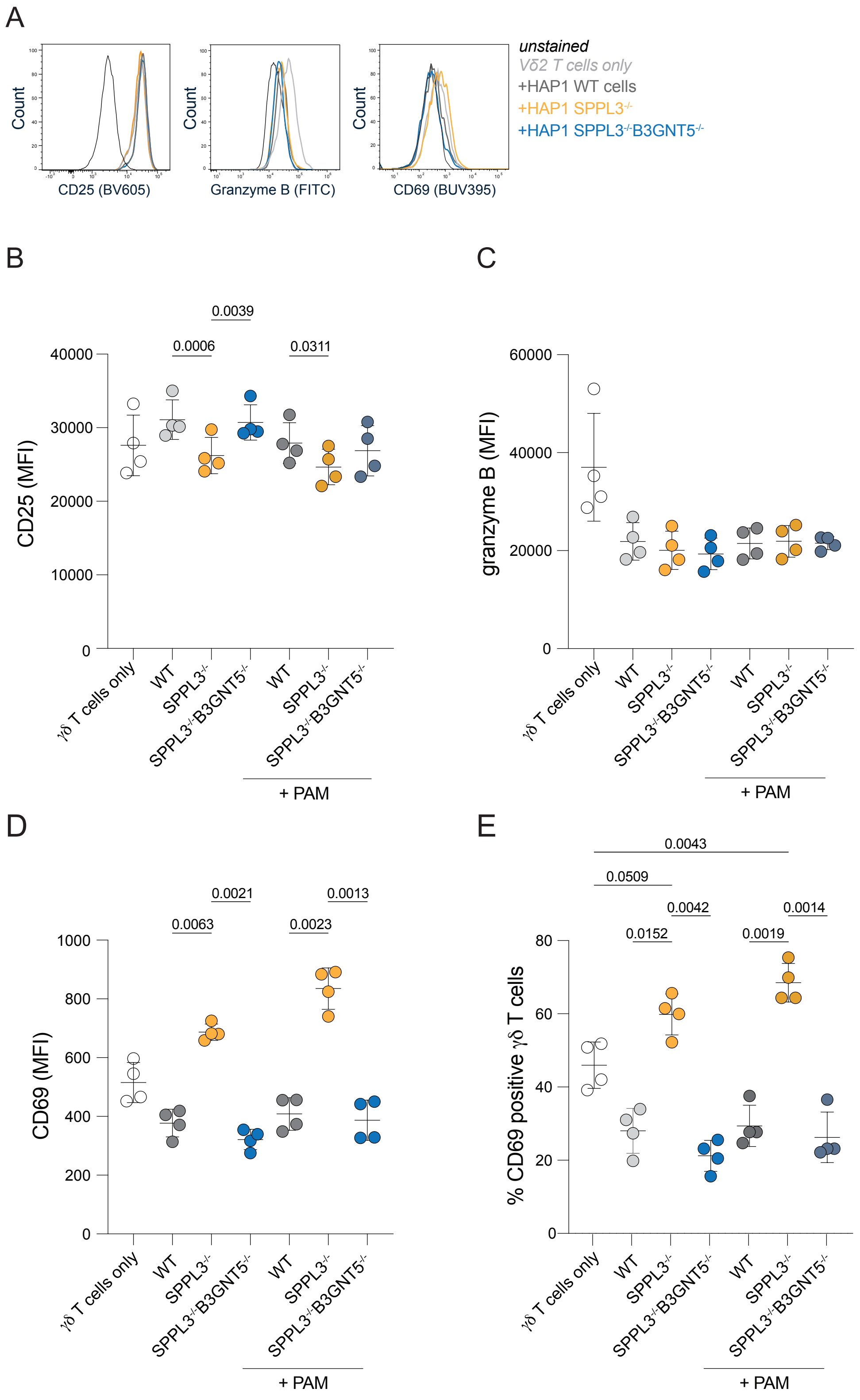
